# Sodium accumulation in breast cancer predicts malignancy and treatment response

**DOI:** 10.1101/2021.04.14.439494

**Authors:** Andrew D James, Theresa K Leslie, Joshua D Kaggie, Laura Wiggins, Lewis Patten, John Murphy O’Duinn, Swen Langer, Marie-Christine Labarthe, Frank Riemer, Gabrielle Baxter, Mary A. McLean, Fiona J Gilbert, Aneurin J Kennerley, William J Brackenbury

## Abstract

Breast cancer is the leading cause of cancer-related death in women worldwide. Development of novel noninvasive diagnostic and predictive pathophysiological biomarkers would represent a significant clinical improvement. Here, we explored the utility of non-invasive ^23^Na MRI to profile tumour physiology using preclinical mouse models of breast cancer. We establish that tissue Na^+^ concentration ([Na^+^]) is elevated vs non-tumour regions across multiple different tumour models. *Ex vivo* SBFI fluorescence imaging corroborated that this elevation in tumour [Na^+^] is due to increased intracellular [Na^+^]. Effective treatment with cytotoxic chemotherapy reduced tumour tissue [Na^+^], but was not detected by ^1^H diffusion-weighted imaging (DWI). Moreover, combining ^23^Na MRI and DWI measurements enabled superior classification accuracy of tumour vs non-tumour regions compared to either parameter alone. Quantification of breast tumour tissue [Na^+^] using ^23^Na MRI thus represents a novel, accurate, non-invasive diagnostic and predictive imaging biomarker.

## Introduction

Metastatic breast cancer represents the leading cause of cancer-related death in women worldwide^1,2^, with the triple-negative subtype associated with particularly poor prognosis^3^. Current standard care pathways involve mammography, ultrasound, magnetic resonance imaging (MRI) and biopsy for diagnostics^4^. The inclusion of additional MRI modalities in care pathways can improve survival outcomes via early diagnosis, structural and morphological assessment of tumours and the monitoring of therapy response^5,6,7^,8. Additional noninvasive MRI biomarkers could therefore contribute a significant clinical improvement.

Recent MR advances have aimed to address the unmet clinical need for better patient stratification. Dynamic contrast-enhanced (DCE) ^1^H MRI has excellent sensitivity for diagnosis and provides functional information about tumour neoangiogenesis^9^. Advances in metabolic imaging, such as the use of PET/MR and hyperpolarized pyruvate (^13^C labelled approaches) have shown promise for measuring breast tumour metabolic activity^10,11^. Nevertheless, these methods are invasive, and are limited by safety concerns surrounding gadolinium-based contrast agents and their requirement for complex equipment, respectively^12^. Alternatively, ^1^H DWI is fully noninvasive, and is used regularly for detecting malignant breast lesions clinically^5,13,14^. DWI determines the local restriction of water within a tissue through quantification of the apparent diffusion coefficient (ADC). Low tissue ADC values are correlated with malignancy in breast cancer^15,16^, and correspond to high cellularity and a smaller extracellular volume fraction. Clinical evidence suggests adjunct DWI can improve the predictive value of conventional DCE MRI^17^.

Previous studies have also indicated that noninvasive ^23^Na MRI of tumour Na^+^ concentration ([Na^+^]) could be a biomarker of malignancy in breast cancer^16,18,19^. The receptivity for ^23^Na MRI is orders of magnitude lower than that of ^1^H MRI^20^ due to the lower tissue concentration of Na^+^ vs H^+^ and an inherently lower gyromagnetic ratio. However, recent technological advances (such as increased magnetic field strength and improved imaging gradients for fast readout strategies) have enabled this pioneering approach to be performed within clinically reasonable timeframes^21^. Indeed, clinical findings indicate that [Na^+^] is elevated within malignant lesions compared with benign lesions^16,18,19^, and findings from two prospective clinical studies suggest that response to neoadjuvant chemotherapy correlates with a decrease in tumour [Na^+^]^19,22^. Moreover, low ADC correlates with elevated total tissue [Na^+^] in malignant lesions^16^, suggesting a link between high cell density and tumour [Na^+^]. As a result, ^23^Na MRI may have utility for breast cancer diagnosis, risk-stratification and monitoring treatment response. However, despite these observations, elevated breast tumour [Na^+^] remains poorly characterised, and urgent supporting data is required for this approach to gain clinical footing and to further understand the underlying pathophysiology.

Identifying the location of Na^+^ accumulation is critically important for determining its pathophysiological impact within the tumour. Increases in total tumour [Na^+^] could reflect changes in intracellular [Na^+^] ([Na^+^]_i_), extracellular [Na^+^] ([Na^+^]_e_), or an increase in the size of the extracellular compartment (extracellular volume fraction, EVF)^23^. Altered [Na^+^]_i_ or [Na^+^]_e_ would have dramatic functional implications for tumour biology; elevations in [Na^+^]_e_ are proinflammatory, inhibit immune cell function^24,25^, and promote resistance to chemotherapy^26^, whereas elevations in [Na^+^]_i_ (reported in cultured cancer cells *in vitro*^,27,28^ and in *ex vivo* tumour samples^29^) could reflect alterations in processes dependent on the inward Na^+^ gradient^23^. Regarding the latter, many common characteristics of invasive tumours (such as cell migration and acidification of the tumour microenvironment) are regulated by this inward Na^+^ transport^27,30^, For example, the Na^+^ dependent Na^+^/H^+^ exchanger (NHE1) regulates tumour pH and metastasis^31^; similarly, NHE1 blockade with cariporide disrupts tumour pH regulation^32^, tumour growth^33^ and sensitises cancer cells to chemotherapy^34^. Moreover, voltage-gated Na^+^ channels (VGSCs) expressed in cancer cells pass a persistent inward Na^+^ current, regulate metastatic cell behaviour and can be inhibited using existing antiepileptic medications^35,36,37,27,38^. Determining the precise location and mechanism of tumour Na^+^ accumulation could therefore reveal novel druggable targets^38^.

In this study, we used mouse models of breast cancer to assess whether the effects of chemotherapy on breast tumour [Na^+^] are reproducible beyond early clinical observations^19,22^ and investigated whether specifically targeting Na^+^ conductance mechanisms affects tumour [Na^+^]. To date, these questions have not been addressed in preclinical *in vivo* models of breast cancer. Our results indicate that multiple orthotopic tumour models (MDA-MB-231, EMT6, 4T1) exhibit elevated tumour [Na^+^], thus recapitulating the clinical picture. Using this preclinical approach, we provide *ex vivo* evidence that the elevated tumour [Na^+^] reflects high [Na^+^]_i_ rather than [Na^+^]_e_. Furthermore, we show that effective treatment with the chemotherapeutic drug docetaxel reduces tumour [Na^+^] but is not reflected by changes in ADC. Importantly, we found that while tumour [Na^+^] has a similar predictive capacity to ADC for classifying malignant regions, combination of these parameters provided superior prediction accuracy. These findings indicate that ^23^Na MRI has utility as both a noninvasive biomarker for malignant disease and treatment response, and position aberrant intracellular Na^+^ handling as a critical feature of malignant breast tumours that may represent an important therapeutic target^23^.

## Results

### ^23^Na MRI reveals elevated tumour [Na^+^] in a preclinical model of triple-negative breast cancer

Xenograft tumours arising from orthotopic implantation of MDA-MB-231 cells represent a model of highly invasive, triple-negative breast cancer. Across a four-week longitudinal study, ^23^Na MRI revealed tissue [Na^+^] is significantly elevated within MDA-MB-231 xenografts compared with contralateral non-tumour regions (imaging performed at Week 2, 3 and 4 post-implant, tumour growth shown in Supplemental Figure 1). ^23^Na signal was linear with [NaCl] (0–100 mM, Supplementary Figure 1b). The observation that [Na^+^] was significantly elevated within tumours was independent of coil setup and readout method. A 3 cm ^23^Na surface coil (B_1_ corrected, ^23^Na 2D gradient echo Cartesian acquisition [2D Cartesian], Figure 1ai) revealed significantly elevated maximum ^23^Na signal (increased 2.5-fold at Week 3, Figure 1aiii) and mean ^23^Na signal (increased 0.9-fold at Week 3, Figure 1aii) in tumour regions compared with non-tumour regions. A dual-tuned linear ^1^H/^23^Na volume coil (^23^Na 2D Cartesian, Figure 1bi) also detected significantly higher mean ^23^Na signal (increased 1.3-fold at Week 3, Figure 1Bii) and maximum ^23^Na signal (increased 1.6-fold at Week 3, Figure 1biii) in tumour regions compared with non-tumour regions. ^23^Na 3D gradient echo spiral out acquisition [3D spiral] with the dual-tuned volume coil (Figure 1ci) also found significant elevations in both mean ^23^Na signal (increased 0.5-fold at Week 3, Figure 1cii) and maximum ^23^Na signal (increased 0.8-fold at Week 3, Figure 1ciii) in tumour regions compared with non-tumour regions. Lower normalised ^23^Na signals from 3D spiral acquisition reflect variable T1 relaxation times between phantom and tissue and a shorter acquisition TR (^23^Na 2D Cartesian, 50 ms; ^23^Na 3D spiral, 10 ms). Taken together, these results confirm elevated tumour [Na^+^] within the MDA-MB-231 preclinical breast cancer model.

**Figure 1:**
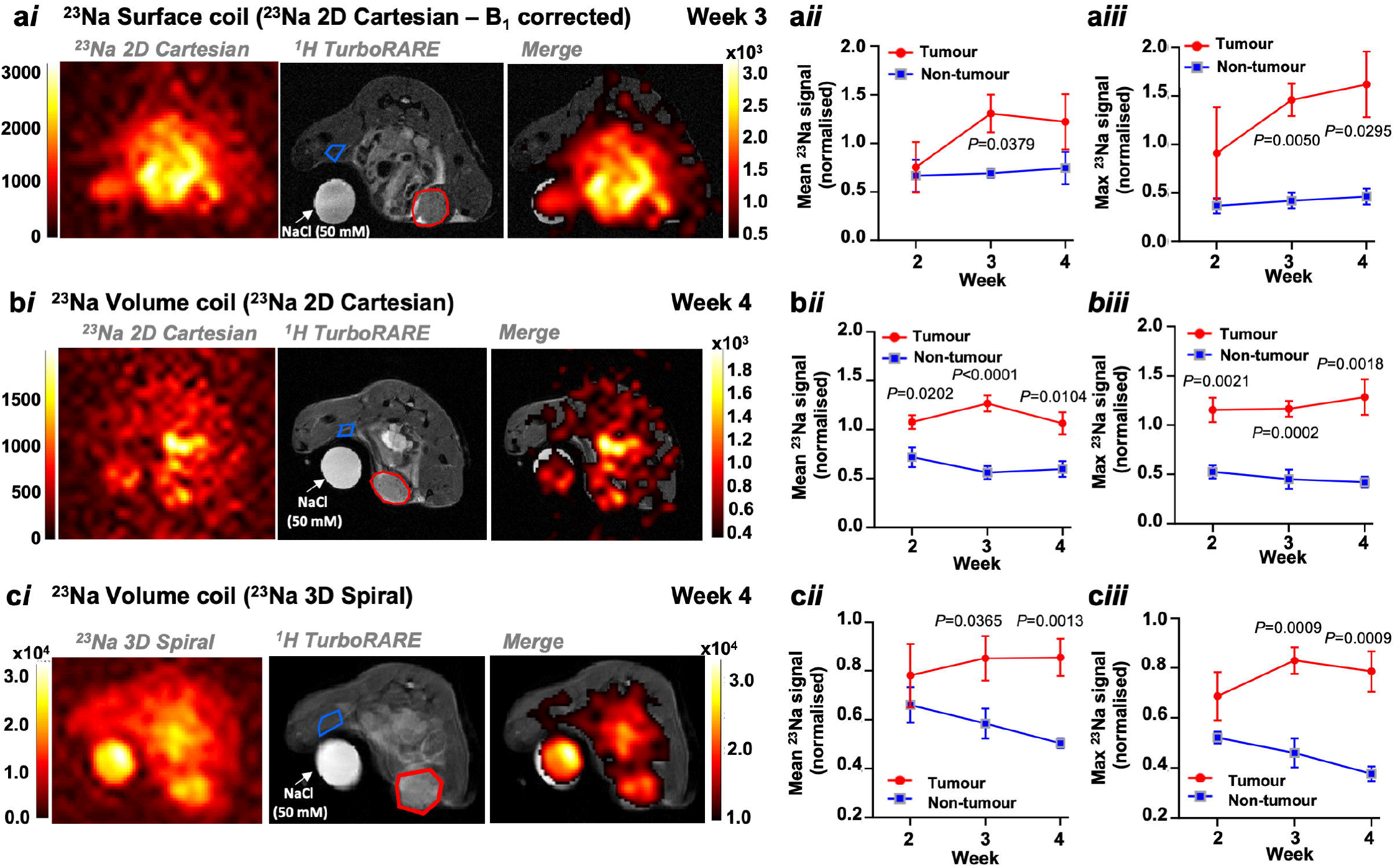
^23^Na MRI reveals elevated [Na^+^] in orthotopic xenograft tumours. ^23^Na imaging was performed in *Rag2*^*−*/−^ *Il2rg*^*−*/−^ mice bearing orthotopic MDA-MB-231 xenograft tumours at 2, 3 and 4 weeks post implantation. Representative images are shown for MRI performed using a bespoke 3 cm ^23^Na surface coil (**a*i***, ^23^Na 2D gradient echo cartesian acquisition (2D cartesian)) or a commercial dual-tuned ^1^H/^23^Na volume coil (**b*i***, ^23^Na 2D cartesian; **c*i***, ^23^Na 3D gradient echo spiral out acquisition (3D spiral)). Regions of interest (ROIs) were defined for both tumour (red region) and for non-tumour contralateral tissue (blue region). Mean and maximum ^23^Na signals from ROIs were normalised to an NaCl phantom of known [Na^+^] (50 mM) to provide a concentration measurement. Mean and maximum ^23^Na signals for tumour vs non-tumour regions are shown for the ^23^Na surface coil (**a*ii*, a*iii***, ^23^Na 2D cartesian, n=3 mice at all timepoints) and dual-tuned ^1^H/^23^Na volume coil (**b*ii*, b*iii*** ^23^Na 2D cartesian. Week 2, n=5; Week 3, n=6; Week 4, n=5. **c*ii*, c*iii*** ^23^Na 3D spiral. Week 2, n=3; Week 3, n=6; Week 4, n=6). Data represent group mean ± SEM. *P*-values for significant differences between groups calculated using an unpaired, two-tailed Student’s t-test.

### ^23^Na MRI improves the predictive potential of ADC for classification of malignant lesions

Low ADC correlates with high cellularity, malignancy^13,39^ and elevated tumour [Na^+^]^16^; ^1^H DWI is used as an adjunct imaging modality for assessing malignant breast lesions in the clinic^14^. To assess whether tumour [Na^+^] is similarly an important biomarker of malignancy, we used machine learning approaches to compare the predictive power of tumour [Na^+^] vs ADC in classifying tumour vs non-tumour regions in the MDA-MB-231 tumour xenograft model. Across all timepoints, ADC was consistently significantly lower in tumour regions than in non-tumour regions (0.7-fold lower at Week 3, Figure 2a, b). Tumour volume was not correlated with either ADC (Figure 2c) or ^23^Na signal (Figure 2d), indicating that both parameters were independent of tumour size. However, In line with previous clinical observations^16^, ADC (all timepoints/regions pooled) was significantly inversely correlated with both maximum ^23^Na signal (r=-0.7499; Figure 2ei) and mean ^23^Na signal (r=-0.6965; Figure 2eii).

**Figure 2:**
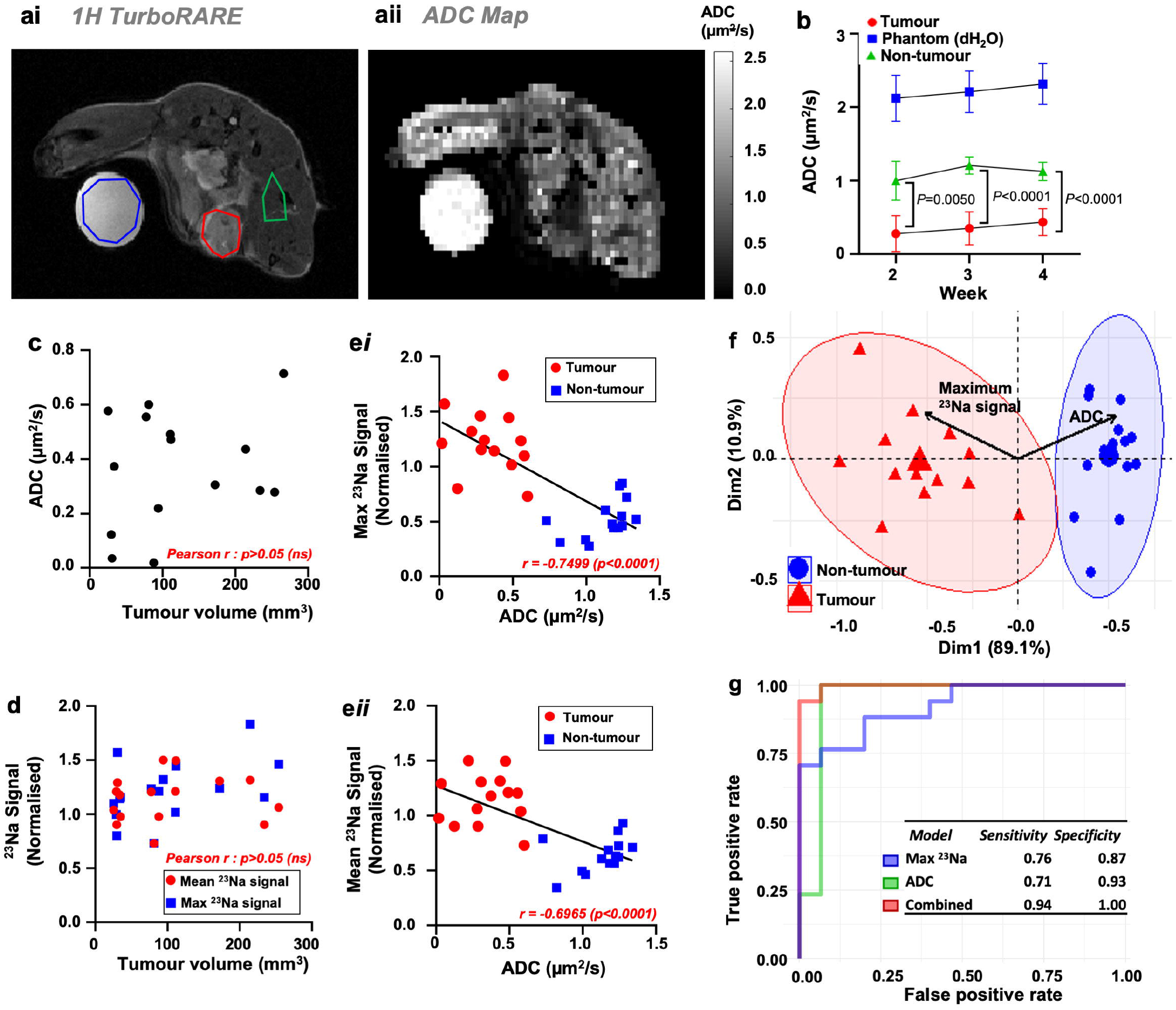
^23^Na MRI combined with and DWI improves tumour vs non-tumour classification accuracy. Corresponding (**a*i***) ^1^H TurboRARE images and (**a*ii***) reconstructed ADC maps (diffusion tensor imaging, b values of 100, 300 and 700 ms) were obtained from mice bearing MDA-MB-231 xenograft tumours at 2, 3 and 4 weeks post implantation. Tumour (red), phantom (blue, 50 mM NaCl) and non-tumour (green) regions of interest (ROIs) are shown. (**b**) ADC in tumour, phantom and non-tumour regions ROIs at each imaging timepoint (Week 2, n=4; Week 3, n=5; Week 4, n=6. mean ± SEM) (**c**) Tumour volume does not correlate with tumour ADC (pooled ROIs across all timepoints, n=15). (**d**) Neither mean nor maximum ^23^Na signal (^23^Na 2D cartesian acquisition) from tumour ROIs correlate with tumour volume (n=16). Both maximum (***ei***) and mean (***eii***) ^23^Na signal are negatively correlated with ADC across all regions (tumour, red; non-tumour, blue, all timepoints pooled, n=30) (**f**) Principal component analysis biplot of maximum ^23^Na signal and ADC (data from **e*i***) with loading vectors for maximum ^23^Na signal and ADC and 95% concentration ellipses for each ROI type. Receiver operating characteristics are shown (**f**) for linear discriminant models trained on maximum ^23^Na values alone, ADC alone, or combination (training data from ***ei***, test data from DTX control cohort (Figure 5, n=32 regions from all timepoints). Classification sensitivity and specificity is presented inset. *P*-values for significant differences between groups calculated using an unpaired, two-tailed Student’s t-test and for correlation using a Pearson r test.

To compare whether mean or maximum ^23^Na signal from tumour and non-tumour regions performed better at classifying regions, we used data from Figure 2ei and 2eii to train two single-variable linear discriminant analysis (LDA, a supervised classification algorithm) models and performed leave-one-out cross validation (LOOCV). Maximum ^23^Na signal achieved a classification accuracy of 92.9%, whereas mean ^23^Na signal achieved 89.3%. Maximum ^23^Na signal was therefore carried forward for our comparisons with ADC. Principal component analysis of maximum ^23^Na signal and ADC reported clear separation of tumour and non-tumour regions (95% concentration ellipses, Figure 2f). The distinct opposing direction and similar size of the associated loading vectors (Figure 2f, black arrows) demonstrate that maximum ^23^Na signal and ADC are inversely correlated yet hold almost equivalent dominance along principal component 1, suggesting that these variables are equally important in explaining the variance between the tissue groups.

To test the comparative predictive capacity of maximum ^23^Na signal vs ADC, we trained LDA models (data from Figure 2ei) and classified novel data taken from an independent experimental cohort (docetaxel control group, see Figure 4). LDA models trained on either maximum ^23^Na signal or ADC alone achieved accuracies of 81.2%, while the model trained on a combination of both parameters achieved an accuracy of 96.9%. Moreover, the maximum ^23^Na signal or ADC single parameter models achieved sensitivities of 0.76 and

0.71 and specificities of 0.87 and 0.93, respectively, whereas the combined parameter model achieved a sensitivity of 0.94 and a specificity of 1.0 (area under the receiver operating characteristics plot, Figure 2g; confusion matrices are in Supplementary Figure 2a). Together, these data indicate that an LDA model trained on maximum tissue [Na^+^] achieves similar classification accuracy of tumour vs non-tumour regions to one trained on tissue ADC. Importantly, however, a combination of these two parameters holds superior predictive power for distinguishing malignant lesions from non-tumour regions.

### Elevated tumour [Na^+^] is due to increased intracellular [Na^+^]

In agreement with the ^23^Na MRI data, inductively-coupled plasma mass spectrometry (ICP-MS) analysis of dissected MDA-MB-231 tumour samples (Figure 3a) showed that total tissue [Na^+^] was significantly higher in xenograft tumours (46.9 ± 5.4 mM) compared with healthy mammary glands from naive mice (29.7 ± 1.9 mM, Figure 3b). Identifying the compartmentalisation of this Na^+^ accumulation is critically important for determining the impact of elevated [Na^+^] on tumour pathophysiology. Because the low tumour ADC indicates a larger intracellular volume fraction compared with non-tumour regions, the elevated tumour [Na^+^] is unlikely to reflect an increase in the extracellular volume fraction. We hypothesised that elevated tumour [Na^+^] instead reflects an increase in intracellular [Na^+^] ([Na^+^]_i_). To test this, we used complementary *ex vivo* approaches to measure [Na^+^]_i_ and extracellular [Na^+^] ([Na^+^]_e_) within acutely isolated live tumour tissue slices.

**Figure 3:**
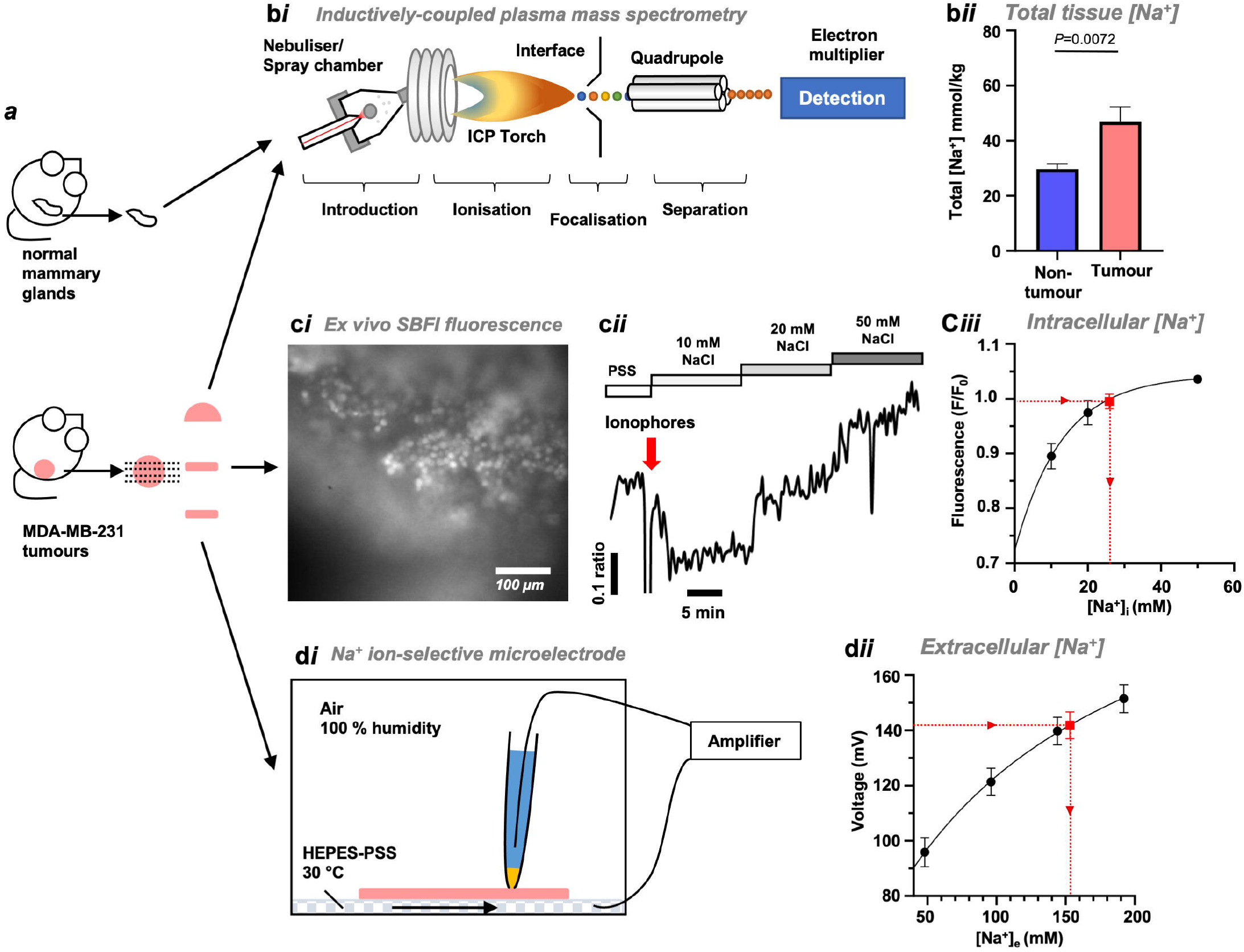
Intracellular [Na^+^] is elevated in live xenograft tumour slices. (**a*i***) Following Week 4 imaging, tumours were acutely isolated, divided and prepared for inductively-coupled plasma mass spectrometry (ICP-MS, **b*i***) analysis or *ex vivo* measurements of [Na^+^]_i_ and [Na^+^]_e_. Healthy mammary glands (HMGs) from naive female mice were included in ICP-MS analysis. (**b*ii***) Total tissue [Na^+^] in MDA-MB-231 xenograft tumours (n=10) and HMGs (n=11). **c*i***, MDA-MB-231 xenograft tumour slices (200 µm) were loaded with SBFI-AM (10 µM) and immobilised for imaging (**c*ii***). Baseline SBFI fluorescence was recorded during perfusion with HEPES-buffered physiological saline solution (PSS) containing 144 mM NaCl. to calibrate [Na^+^]i for each cell *in situ*, perfusion was switched to 10 mM NaCl, followed by application of the ionophores (red arrow) gramicidin (3 µM), monensin (10 µM) and ouabain (1 mM). Tumour slices were sequentially perfused with 20 mM and 50 mM NaCl. The final three frames acquired during each phase were used to determine average fluorescence for each calibration step (**c*iii***) Intracellular [Na^+^] during the baseline PSS phase (red point) was calibrated (n=3 slices from 2 separate animals, 11–16 cells per experiment, F/F_0_) using nonlinear regression (one-phase association). (**d**) Concurrently, Na^+^ ion selective microelectrodes (ISMEs) calibrated with PSS of variable [Na^+^] (48 mM, 96 mM, 144 mM, 192 mM, no replacement ion) were used to measure extracellular [Na^+^] (red point) in MDA-MB-231 xenograft tumour slices (500 µm, n=6 slices from 6 separate animals, 12 recordings per slice). Recordings were interpolated using nonlinear regression (Padé (1,1,) approximant). Data represent group mean ± SEM.

**Figure 4:**
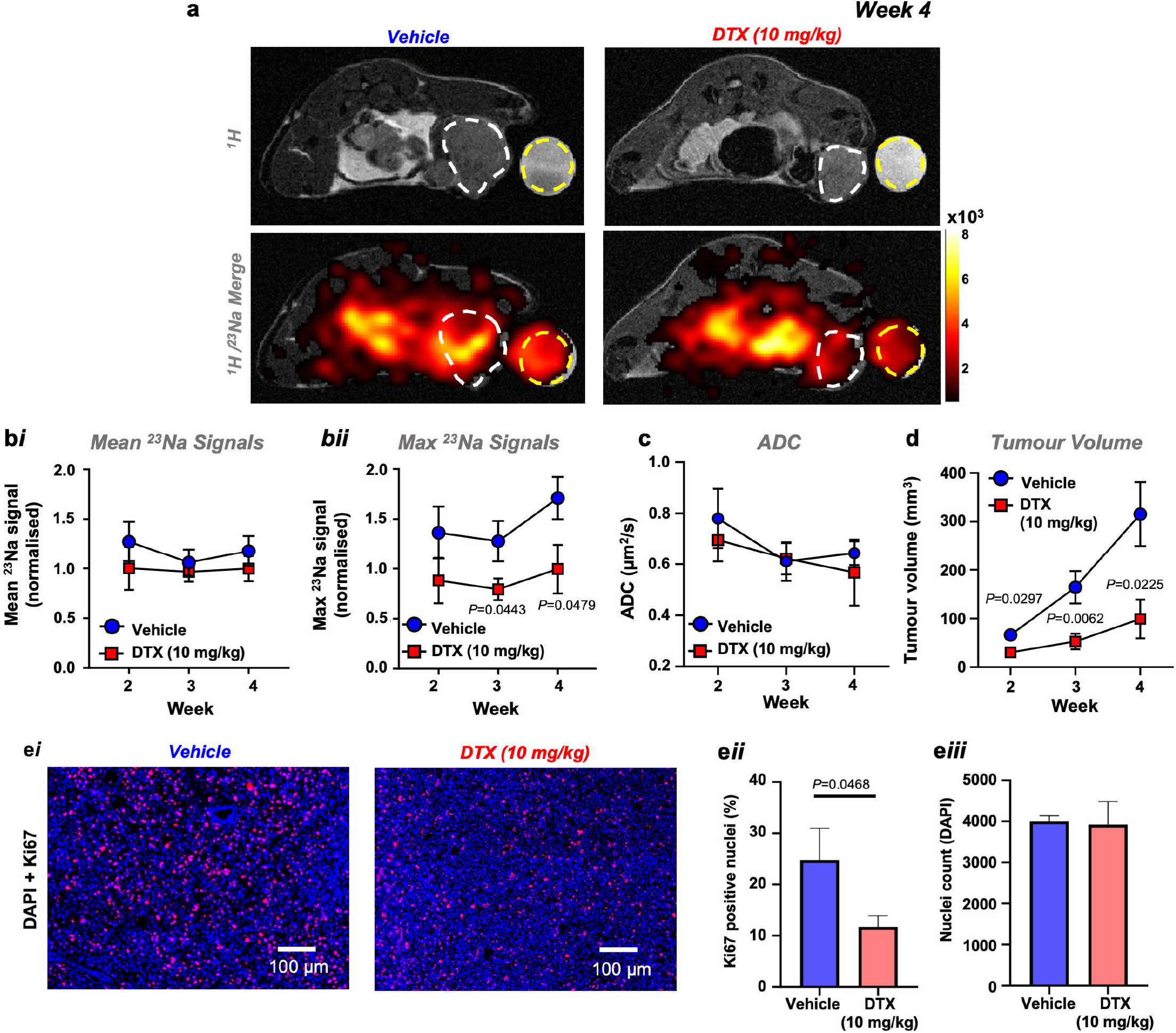
Docetaxel decreases maximum ^23^Na signal detected within MDA-MB-231 xenograft tumours. Following orthotopic MDA-MB-231 cell implantation, *Rag2*^*−*/−^ *Il2rg*^*−*/−^ mice were randomised to receive either docetaxel (10 mg/kg) or vehicle i.p. once weekly. (**a**) Representative ^1^H TurboRARE and ^1^H/^23^Na 2D cartesian (merge) images for docetaxel and vehicle treated mice at Week 4 post implant; regions of interest (ROIs) for tumour (white dash) and NaCl phantom (50 mM, yellow dash) are annotated. Both mean (**b*i***) and maximum (**<u>b</u>*<u>ii</u>***) ^23^Na signals measured from tumour ROI at Weeks 2, 3 and 4 post xenograft implantation (normalised to phantom to provide a concentration measurement. Vehicle, Week 2, n=7; Week 3, n=6; Week 4, n=7. Docetaxel, Week 2, n=9; Week 3, n=8; Week 4, n=7). (**c**) Effect of docetaxel on tumour ADC (Vehicle, Week 2, n=5; Week 3, n=6; Week 4, n=7. Docetaxel, Week 2, n=6; Week 3, n=8; Week 4, n=7). (**d**) Effect of docetaxel treatment on tumour growth rate (vehicle, n=6 for all timepoints. Docetaxel, Week 2, n=8; Week 3, n=8; Week 4, n=5. Volume measured by multislice (1 mm) ^1^H TurboRARE imaging. (**e*i***) Representative images of Ki67/DAPI stained tissue tumour sections, with (**e*i***) quantified Ki67 positive nuclei (vehicle, n=5; docetaxel, n=7; 2–10 regions each section) and (**e*ii***) nuclei counts (vehicle, n=5; docetaxel, n=6; 1–10 regions each section). Data represent group mean ± SEM. *P*-values for significant differences between groups calculated using an unpaired, two-tailed Student’s t-test (**bi, bii, c, d**) or an unpaired, two-tailed nested t-test (**eii, eiii**).

Using the Na^+^ selective dye SBFI-AM (10 µM) and an *in situ* calibration with dynamic range comparable to that previously described^40^ (Figure 3cii), we found that resting [Na^+^]_i_ in acutely isolated MDA-MB-231 tumour slices (Figure 3ci, isolated following Week 4 imaging timepoint) was 25.9 ± 1.0 mM (Figure 3ciii). This is higher than the typical [Na^+^]_i_ for healthy mammalian cells (5–15 mM)^41^ and yet is in line with other *in vitro* fluorescence studies showing high [Na^+^]_i_ in cultured cancer cell lines^28,42,43^. In contrast, Na^+^-selective microelectrodes^44^ (Figure 3di) showed that the [Na^+^]_e_ within slices from the same tumours was 157.8 ± 1.0 mM (Figure 3dii). This is within the expected serum Na^+^ concentration range for female mice of BALBc lineage^45^. Taken together with the DWI data, these results indicate that elevated tumour [Na^+^] is due to an increased [Na^+^]_i_ whereas [Na^+^]_e_ is unchanged.

### Detection of docetaxel treatment response by ^23^Na MRI in MDA-MB-231 xenografts

Using a randomised, controlled, interventional study approach, we sought to determine whether treatment response can be detected using ^23^Na MRI in xenograft tumour bearing mice. There is initial evidence of this within two prospective clinical studies with small cohort sizes (n=15^19^ and n=5^22^ patients) which found that elevated [Na^+^] in malignant breast lesions was reduced in patients responding to neoadjuvant chemotherapy.

Following tumour cell (MDA-MB-231) implantation, mice were treated with docetaxel 10 mg/kg i.p.) or vehicle once weekly from Week 1 post implantation. Docetaxel treatment significantly inhibited tumour growth; at Week 3, tumour volume within the treatment group was 32.2% of that within the vehicle group (Figure 4d). Furthermore, docetaxel treatment significantly reduced Ki67-positive nuclei within tumour sections by 52.7% (Figure 4ei and Figure 4eii) despite no change in nuclei count (Figure 4eiii). Moreover, docetaxel treatment led to a significantly lower maximum ^23^Na signal within tumour regions compared with untreated control mice (37.8% lower at Week 3, Figure 4ai, bii). This decrease was statistically significant at the Week 3 and Week 4 timepoints, suggesting that localised changes in tumour [Na^+^] reflects clinical response. However, no difference was observed between docetaxel and control groups for either mean tumour ^23^Na signal (Figure 4bi) or isolated tissue sample [Na^+^] (ICP-MS, Supplementary Figure 3a). The difference between maximum and mean ^23^Na signals potentially reflects important intratumoural heterogeneity requiring improved MRI resolution to probe further.

Previous studies indicate that low ADC correlates with high expression of the proliferation marker Ki67^46–48^. Given the decrease in maximum ^23^Na signal observed upon treatment with docetaxel, we performed DWI to assess whether treatment response was reflected in ADC. In agreement with the tumour section nuclei counts, there was no difference in ADC between control and docetaxel groups (Figure 4c). These data indicate that the docetaxel-induced change in tumour [Na^+^] was not due to altered cellularity, suggesting that tumour [Na^+^] might provide improved treatment monitoring over ADC.

### Effect of cariporide and eslicarbazepine acetate on tumour [Na^+^]

Na^+^ channels and transporters utilise the inward electrochemical Na^+^ gradient to achieve their physiological function, with many implicated in tumour progression and metastasis^23^. The elevated [Na^+^]_i_ in intact tumour slices could reflect increased Na^+^ transporter activity. We therefore investigated whether inhibition of either NHE1 (cariporide) or VGSCs (eslicarbazepine acetate, ESL) affected elevated [Na^+^] within MDA-MB-231 xenograft tumours. ESL and its major active metabolite (S)-licarbazepine^49^ inhibit inward Na^+^ current in MDA-MB-231 cells^50^, and was selected due to its preferred use in the clinic compared to older VGSC-inhibiting antiepileptic medications such as phenytoin^51^. Moreover, cariporide was administered at a dose that previously exhibited efficacy in *in vivo* models of cancer^32,33,34^.

ESL (200 mg/kg daily p.o. from day 7 post implant) had no effect on mean (Figure 5a, 5bi) or maximum ^23^Na signal (Figure 5a, 5bii) within the tumour compared with untreated control mice. Similarly, ESL had no effect on tumour volume (Figure 5c). Cariporide (3 mg/kg daily i.p.^33,34^ from day 7 post implant) also did not reduce mean or maximum ^23^Na signal within the tumour compared with untreated control mice; in fact there was a significant increase in mean ^23^Na signal at Week 2 and maximum ^23^Na signal at Week 3 (0.55-and 0.44-fold, respectively; Figure 5d,e). Nevertheless, cariporide had no effect on tumour growth (Figure 5f). In agreement with these observations, there was no effect of ESL or cariporide on VGSC gating properties or peak current density measured in acutely isolated tumour slices (Supplementary Figure 4). This was despite achieving plasma [licarbazepine] (1 hour post dosing) comparable to that showing antiepileptic efficacy in the clinic^52^ (plasma, 6.5 ± 0.9 ng/µl; tumour, 4.5 ± 0.8 ng/mg, n=3 tumours). Taken together, these data indicate that ESL and cariporide at the current doses elicit no detectable decrease in tumour [Na^+^], and suggest that alternative (or multiple) Na^+^ transporters should be targeted to alter elevated tumour [Na^+^] *in vivo*.

**Figure 5:**
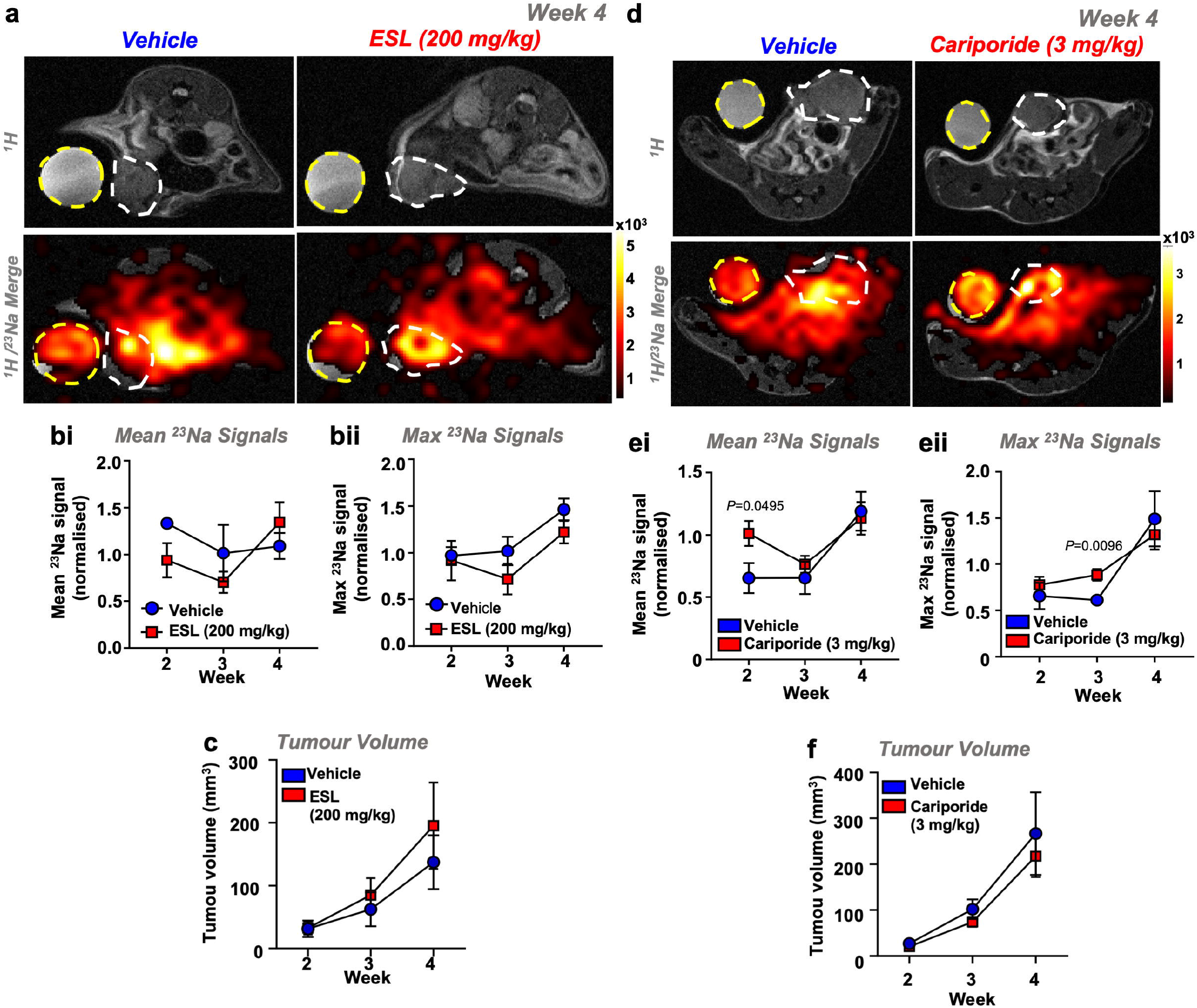
Cariporide and eslicarbazepine acetate have no effect on tumour [Na^+^] in MDA-MB-231 xenografts. *Rag2*^*−*/−^ *Il2rg*^*−*/−^ mice bearing MDA-MB-231 tumours were randomised to receive either cariporide (3 mg/kg) vs vehicle i.p. once daily or eslicarbazepine acetate (ESL, 200 mg/kg) vs vehicle p.o. once daily commencing 1 week post xenograft implantation. Representative ^1^H TurboRARE and ^1^H/^23^Na 2D cartesian (merge) images for (***a***) cariporide vs vehicle (***d***) ESL vs vehicle at Week 4 post implant; regions of interest (ROIs) for tumour (white dash) and NaCl phantom (50 mM, yellow dash) are annotated. Mean (***bi, ei***) and maximum (***bii, eii***) ^23^Na signals from tumour ROIs (normalised to phantom to provide a concentration measurement) were measured at Weeks 2, 3 and 4 post xenograft implantation (cariporide group: n=5 per group per timepoint. ESL vehicle group: Week 2, n=3; Week 3, n=3; Week 4, n=4. ESL treatment group: Week 2, n=4; Week 3, n=4; Week 4, n=5). Tumour growth rate was measured by multislice ^1^H TurboRARE imaging (**c**, cariporide. Vehicle Week 4, n=4; all other data points n=5. **f**, ESL. Vehicle, n=4; treatment group, n=5). Data represent group mean ± SEM. *P*-values for significant differences between groups calculated using an unpaired, two-tailed Student’s t-test.

### Tumour [Na^+^] is elevated across multiple murine models of breast cancer

To confirm the clinical relevance of these findings, we next assessed whether elevated tumour [Na^+^] is a common feature across other orthotopic murine models of breast cancer. We selected the EMT6 (Figure 6a) and 4T1 (Figure 6b) murine mammary carcinoma cell lines, which form solid tumours in immunocompetent BALB/c mice. These cell lines give rise to luminal A tumours that are weakly ER-positive, claudin-low (EMT6) and ER-negative and non-claudin low (4T1). Moreover, these models exhibit significant immunogenicity and leukocytic infiltration, with EMT6-derived tumours exhibiting higher GR1+ granulocyte and CD3+ T-cell infiltration^53^.

**Figure 6:**
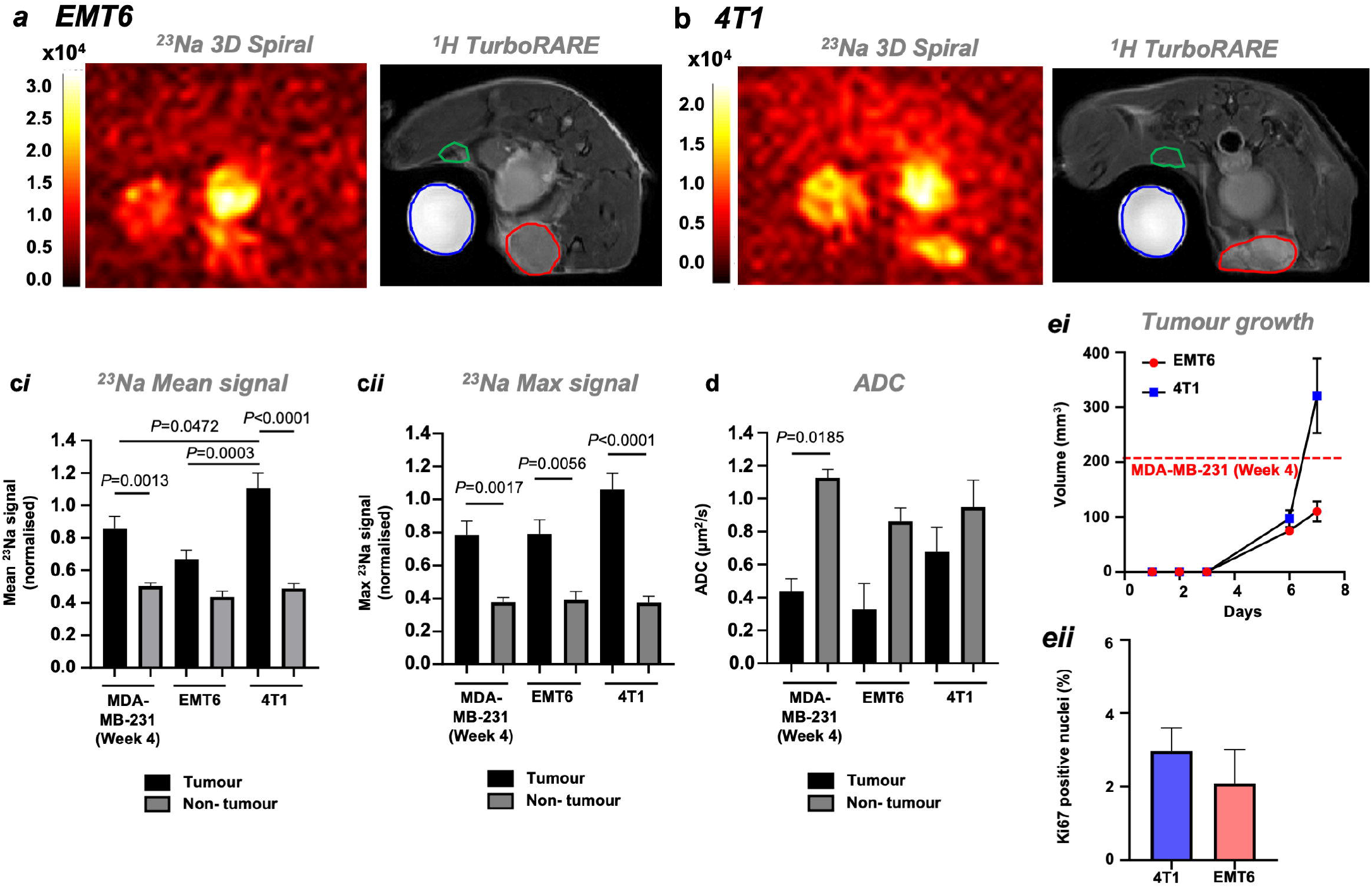
Elevated tumour [Na^+^] is a common feature of preclinical models of breast cancer. Representative ^23^Na 3D gradient echo spiral out and ^1^H TurboRARE images are shown for orthotopic (**a**) EMT6 or (**b**) 4T1 allograft tumours, imaged at day 7 post-implantation (tumour region of interest (ROI), red; non-tumour ROI, blue). Mean ^23^Na signal (**c*i***), maximum ^23^Na signal (***Cii***) within tumour and non-tumour ROIs were normalised to NaCl phantom (50 mM) to provide a concentration measurement; data from MDA-MB-231 tumours at Week 4 post implantation included for statistical comparison. ADC (**d**) was assessed by DWI. (**e*i***) Tumour volume (4T1 and EMT6) was calculated using the modified ellipsoidal formula *volume = 1/2(length × width*^*2*^); dotted line denotes the average size of MDA-MB-231 tumours at 4 weeks post implantation (see Supplemental Figure 1A). (**e*i***) quantified Ki67 positive nuclei for 4T1 and EMT6 tumour sections (both groups n=4, 3–12 regions each section). Data represent group mean ± SEM (4T1, EMT6, n=5; MDA-MB-231, n=6). *P*-values for significant differences between groups calculated using a one-way ANOVA with post-hoc Tukey test for multiple comparisons (**c*i*, c*ii***), a Kruskal-Wallis test with post-hoc Dunn’s test for multiple comparisons (**d**) or a nested t-test (**e*ii***).

EMT6 and 4T1 tumours grew rapidly after implantation (Figure 6ei), reaching a similar size at 1 week post implant to MDA-MB-231 cells at 4 weeks post implant; no difference was observed in Ki67 density between the two models (Figure 6eii). At 1 week post implant, ^23^Na MRI revealed elevated [Na^+^] in both EMT6 (Figure 6a) and 4T1 (Figure 6b) tumours. In the 4T1 model, mean ^23^Na signal was significantly elevated compared with non-tumour regions (0.5-fold increase, Figure 6ci). Similarly, maximum ^23^Na signal was significantly elevated (1-fold increase, EMT6; 1.8-fold increase, 4T1; Figure 6cii) within tumour regions compared with non-tumour regions. Interestingly, 4T1 tumours exhibited significantly higher mean ^23^Na signal compared with EMT6 or MDA-MB-231 tumours (0.7-fold and 0.2-fold increases, respectively; Figure 6ci). These results suggest that although elevated tumour [Na^+^] is consistent across multiple models of breast cancer, the absolute tumour [Na^+^] is variable, hinting at important underlying biological differences in Na^+^ handling across these tumour types. However, in contrast to the MDA-MB-231 model, no significant difference was observed in ADC between tumour and non-tumour regions in EMT6 and 4T1 models (Figure 6d). Similar to [Na^+^], these data suggest that tumour ADC is variable across these models. Importantly, however, given that elevated tumour [Na^+^] was identified by ^23^Na MRI in the 4T1 model but no difference was seen in ADC, ^23^Na MRI might in certain cases be better positioned than DWI to discern malignancy.

## Discussion

The present study is the first to demonstrate that tumour [Na^+^] is elevated relative to non-tumour regions in the orthotopic MDA-MB-231 xenograft and 4T1 and EMT6 allograft models of breast cancer. Moreover, while supervised classification algorithms trained on ^23^Na MRI data achieved similar accuracy to those trained on DWI data (ADC) in distinguishing tumour vs non-tumour regions, a model trained on both of these parameters combined achieved superior classification accuracy, suggesting that ^23^Na MRI holds promise as a novel diagnostic readout. Using a randomised, controlled study approach, we show that docetaxel, an antineoplastic drug used clinically that disrupts microtubule formation and cell division, reduces maximum tumour [Na^+^] while slowing tumour growth rate but has no effect on ADC; these findings suggest that ^23^Na MRI also has utility as a readout of response to chemotherapy, and may be better suited for this purpose than DWI. These findings broadly confirm for the first time in a randomised, controlled, interventional study design, the chemotherapy response effect reported in previous small-cohort clinical observational studies^18,19^. We show in intact breast tumour slices that elevated tumour [Na^+^] reflects increases in [Na^+^]_i_, while extracellular [Na^+^]_e_ is unaltered. This aberrant intracellular Na^+^ handling sheds light on a previously underappreciated characteristic of malignant breast lesions, and has considerable implications for pathophysiological processes linking ion homeostasis to tumour cell function.

The finding that tumour [Na^+^] is elevated within mouse models of breast cancer is consistent with previous observations from clinical studies^16,18,19^. Ouwerkerk et al. reported that mean total tissue [Na^+^] was ∼53 mM (n=19 patients) in malignant lesions and ∼26 mM in benign lesions (n=3 patients)^18^. Similarly, Zaric et al. reported a mean total tissue [Na^+^] of ∼69 mM in malignant lesions compared with ∼35 mM in normal glandular breast tissue (n=17 patients)^16^. These previously reported values are broadly in agreement with our ICP-MS and ^23^Na MRI data. However, our finding that maximum ^23^Na signal improves distinction of tumour vs non-tumour regions compared to mean ^23^Na signal potentially reflects considerable heterogeneity of [Na^+^] within tumours. Indeed, despite limited spatial resolution, we observed clear variability in ^23^Na signal across the tumour regions (Figures 1, 4, 6). This heterogeneity could also explain why docetaxel treatment conferred a reduced maximum ^23^Na signal within tumours despite no difference in mean ^23^Na signal. This has broader implications for prognosis, since intratumoural heterogeneity is considered a key contributor to resistance to chemotherapy^54^, and suggests a potential link between local changes in tumour [Na^+^] and cancer cell function that could be discerned using ^23^Na MRI. Realisation of improved spatial specificity for ^23^Na MRI would enable incorporation of tumour [Na^+^] heterogeneity within more complex analyses. Indeed, a recent study coregistering MDA-MB-231 and 4T1 tumour section histology with various ^1^H parameters (including DCE MRI and DWI) found that DCE MRI could distinguish tumour subregions (i.e. hypoxic vs normoxic and viable vs nonviable regions)^55^. Inclusion of (or replacement with) non-invasive ^23^Na readouts would offer an important additional dimension to such functional imaging methods.

The present study also establishes that tumour [Na^+^] coregisters with a lower ADC within preclinical models of breast cancer and that these parameters are inversely correlated across non-cancerous and malignant lesions, recapitulating previous clinical observations^13,16,39,56^. These findings suggest a functional link between high cellularity and Na^+^ accumulation in tumours, supporting the notion that elevated tumour [Na^+^] is likely due to changes within the intracellular compartment. Interestingly, principal component analysis revealed that ADC and maximum ^23^Na signal held almost equivalent importance when explaining variance across all regions (tumour and non-tumour). Linear discriminant analysis models trained on either ^23^Na signal or ADC were comparable in their ability to correctly classify regions. However, a model trained on both variables achieved superior classification accuracy. One can therefore hypothesise that their predictive resolutions do not completely overlap (see Supplemental Figure 2) and are indeed complementary.

While recent recommendations have been published^57^, there is as yet no clearly established consensus on optimal DWI methodology or the threshold ADC value with which to denote malignancy^58^. Moreover, DWI sensitivity and ADC quantification are influenced by differences in methodology (i.e. choice of b-values, magnet strength) and tumour heterogeneity^59,60,61^, thereby preventing the establishment of a generalised threshold ADC value. Indeed, care must be taken when inferring cellularity and malignancy from DWI, as ADC negatively correlates well with cellularity in some tumours but not in others^62^. Taken together with the improved statistical discrimination of 4T1 and EMT6 tumours by ^23^Na MRI compared to DWI, our classification results suggest that a combined approach using both DWI and ^23^Na MRI could improve specificity and predictive accuracy in the clinic, thus enabling superior noninvasive diagnostic and treatment monitoring capabilities beyond DWI alone.

The present study is the first to report an effect of docetaxel on tumour [Na^+^] in the preclinical setting. Previous clinical observations suggested that treatment with chemotherapy lowers tumour [Na^+^] in breast cancer. Two small prospective clinical studies measured significant reductions (27%, n=15 patients^19^; 21%, n=5 patients^22^) in total tissue [Na^+^] using ^23^Na MRI following response to neoadjuvant chemotherapy. In the present study, docetaxel treatment led to a decrease in xenograft tumour growth and a lower maximum tumour ^23^Na signal compared with that in vehicle treated mice (37.8% lower at Week 3). However, the mechanism underlying this effect is not yet known. Similarly, it is not known whether this decrease in tumour [Na^+^] contributes to, or is a consequence of, treatment response. Furthermore, treatment with docetaxel did not decrease ADC, suggesting a mechanism other than changes in cellularity such as decreased tumour vascularity (docetaxel is antiangiogenic)^63^. Moreover, these data suggest that in certain circumstances ^23^Na MRI may be better positioned to discern treatment response than DWI.

Beyond breast cancer, contrasting observations have been made in fibrosarcoma and gliosarcoma models following treatment with cyclophosphamide or carmustine treatment, respectively^64^,65,65,66 In these models, tumour ^23^Na signal increased with tumour growth, and the changes in ^23^Na signal and ADC induced by chemotherapy were correlated^67^. The authors concluded that the correlated changes in ^23^Na and ADC reported the same underlying physiological change, namely decreased cellularity and an increase in the extracellular volume fraction (EVF). This is in contrast to the present study in orthotopic breast tumours, where tumour ADC was unchanged following docetaxel treatment, despite a concomitant decrease in the max ^23^Na signal. These conflicting results indicate that tumour [Na^+^] responses likely differ depending on cancer type (i.e. breast vs gliosarcoma) and the choice of chemotherapeutic intervention.

The present study provides direct evidence from intact, live MDA-MB-231 tumour slices that elevated tumour [Na^+^] arises due to an intracellular accumulation of Na^+^. [Na^+^]_i_ was 25.9 mM within the tumour cells (assessed by SBFI fluorescence), well in excess of the expected upper physiological bound for [Na^+^]_i_ (∼15 mM^41^). This is in agreement with our DWI results, which suggest that extracellular volume fraction is decreased in tumours vs non-tumour regions, and that elevated tumour [Na^+^] must therefore reflect an increase in either [Na^+^]_i_ or [Na^+^]_e_. Moreover, [Na^+^]_e_ was within the expected physical range for female BALB/c mouse serum (∼157–160 mM^45^), indicating that elevated tumour [Na^+^] is not due to changes in the extracellular compartment. Interestingly, the tumour [Na^+^]_i_ we observed *ex vivo* is similar to that extrapolated (∼30 mM) in two patients with triple negative breast cancer using complementary ^23^Na and ^1^H DCE MRI approaches to determine tumour [Na^+^], EVF and water fraction^68^. Furthermore, although at the limit of statistical distinction, intracellular-weighted ^23^Na imaging has found that [Na^+^]_i_ is elevated in prostate peripheral zone tumours compared with adjacent healthy peripheral zone^69^. Given the small size of MDA-MB-231 xenograft tumours, a similar assessment of intracellular Na^+^ by ^23^Na MRI could be enabled by improvements to ^23^Na contrast to noise ratio that are subsequently sacrificed to increase spatial resolution^70^.

Using a simplified model, the relative EVF to intracellular volume fraction (IVF) can be estimated from the values obtained from the present study. If we consider that total tissue [Na^+^] = (1–EVF) x [Na^+^]_i_ + EVF x [Na^+^]_e_, where EVF = 1–IVF^18^, EVF (and thus IVF) can be resolved for given values of [Na^+^]_i_, [Na^+^]_e_ and total tissue [Na^+^]. If we consider [Na^+^]_i_ and [Na^+^]_e_ values recorded in tumour slices (25.9 mM and 157.8 mM, respectively) and the observed total tumour [Na^+^] (46.9 mM), we would expect an EVF of 0.159 and IVF of 0.841 in the tumour. Keeping these EVF and IVF values constant and rearranging the above equation to solve [Na^+^]_i_ for healthy mammary glands (total tissue [Na^+^] = 29.7 mM) with an assumed [Na^+^]_e_ of 158 mM^45^ returns a value of 5.4 mM. This is within the expected physiological range^41^; however, this simplified model does not take into account changes to the EVF:IVF ratio in tumours relative to healthy tissue, nor the blood or ductal compartments. Using shift reagents, one can quantify the contribution of these compartments with magnetic resonance spectroscopic imaging^71,72^, however this would be invasive and therefore of limited clinical use.

This elevated [Na^+^]_i_ has important implications for cancer cell function, as it would likely impact upon processes governed by Na^+^-linked channels and transporters^23^. For example, the Na^+^/K^+^ ATPase is the predominant consumer of intracellular ATP^73^ and its activity increases across the [Na^+^]_i_ physiological range (5–15 mM)^74^, suggesting full activation in cancer cells exhibiting a [Na^+^]_i_ of ∼26 mM. This has important implications for aberrant glycolytic metabolism linked with invasive behaviour^75^, given that plasma membrane cation pumps are largely fuelled by glycolytic ATP^76,77,78^. Similarly, mitochondrial Na^+^/Li^3+^/Ca^2+^ exchanger activation by elevated [Na^+^]_i_ (K_m_ ∼12 mM)^79^ would be expected to alter mitochondrial function and bioenergetics^80,81^. Processes altered by elevated [Na^+^]_i_ could therefore represent novel therapeutic loci.

On the other hand, the elevated [Na^+^]_i_ could reflect increased inward Na^+^ channel or transporter activity^23^. For example, cell volume, pH regulation and migration are regulated by the activity of NHE1^82,83,31^, and elevations in [Na^+^]_i_ via aberrantly expressed VGSCs have been directly linked to invasion and metastasis in breast cancer^27,38,84^. Plasma membrane Na^+^ channels and transporters are attractive pharmacological targets as they are easily accessible to small molecule inhibitors and are targeted by drugs already used in the clinic. However, in contrast to docetaxel, selective inhibition of either NHE1 or VGSCs (with cariporide or ESL, respectively) did not reduce elevated tumour [Na^+^] in the present study at clinically relevant doses. On the other hand, we saw no effect of docetaxel treatment on VGSC gating properties or peak current density in *ex vivo* tumour slices, despite a decrease in maximum tumour ^23^Na signal (evidence indicates that taxanes modulate VGSC activity in cultured MDA-MB-231 cells^85^). Although we cannot rule out subtle, localised changes in [Na^+^] beyond the sensitivity of the present MRI approach, these results suggest that alternative Na^+^-dependent transporters (or the combined effect of several transporters) may be responsible for increased [Na^+^]_i_ in tumours^23^.

In conclusion, this pioneering study highlights ^23^Na MRI as a novel imaging biomarker for breast cancer diagnosis and response to chemotherapy. In addition, we elevate the importance of altered intracellular [Na^+^] handling in breast tumour pathophysiology, and identify altered tumour [Na^+^] as a common characteristic of preclinical breast cancer models. The inclusion of noninvasive ^23^Na MRI in the patient care pathway may therefore represent an important future clinical refinement in the fight against breast cancer. Supported by these results, the future development of noninvasive ^23^Na MRI with improved contrast to noise ratio^70^ (sacrificed to increase spatial resolution) will further empower its use for diagnostics and exploring tumour heterogeneity and function.

## Methods

### Ethics statement

Approval for all animal procedures was granted by the University of York Animal Welfare and Ethical Review Body. All procedures were carried out under authority of a UK Home Office Project Licence and associated Personal Licences.

### Cell culture

MDA-MB-231 cells, a gift from M. Djamgoz (Imperial College London), were cultured in Dulbecco’s modified Eagle’s medium (DMEM 219690-35, Thermo Fisher Scientific) supplemented with 5% foetal bovine serum (FBS) and 4 mM L-glutamine, and were authenticated by short tandem repeat analysis. EMT6 cells (European Collection of Authenticated Cell Cultures) were cultured in Minimum Essential Medium Eagle’s containing Earle’s Balanced Salt Solution (EMEM M2279, Sigma Aldrich, UK) supplemented with 2 mM Glutamine,1% Non Essential Amino Acids (NEAA) and 10% FBS. 4T1 cells were a gift from Dr. Mihaela Lorger (University of Leeds UK, sourced from ATCC), and were cultured in EMEM supplemented with 2 mM L-glutamine, HyClone™ Vitamin Solution (Cytiva, 1x final working concentration), 1% NEAA, 1 mM sodium pyruvate and 10% FBS. All cells were cultured in a humidified atmosphere of air/CO2 (95:5%) at 37 °C and routinely tested for mycoplasma using the DAPI method.

### Orthotopic breast tumour model

At > 6 weeks of age, *Rag2*^*-*/-^ *Il2rg*^*-*/-^ (bred in-house, Balb/c background strain) or Balb/c (Charles River Laboratories, UK) mice were anaesthetised (2% isoflurane in oxygen (2 l/min)) and 5 × 10^5^ MDA-MB-231 (*Rag2*^*-*/-^ *Il2rg*^*-*/-^), 1 × 10^5^ EMT6 or 1 × 10^5^ EMT6 (BALB/c) cells (suspended in Matrigel, 50% v/v in saline, 50 µl volume) were injected into the left inguinal mammary fat pad. Animal weight and condition were monitored daily; tumour growth was monitored using daily calliper measurement. Where possible, tumour volume was calculated from weekly multi-slice ^1^H scans; otherwise, tumour volumes were calculated from calliper measurements using the modified ellipsoidal formula *volume = 1/2(length × width*^*2*^) ^86^. Mice were euthanized at <5 weeks post-implantation or when tumours reached 15 mm diameter and tumours isolated by dissection.

### Drug treatment interventional studies

To assess the effects of docetaxel, cariporide or ESL on tumour ^23^Na signal, mice bearing orthotopic MDA-MB-231 tumours were randomised to receive either vehicle or drug treatment (docetaxel 10 mg/kg in 1:1:20 ethanol:Tween 20:5% glucose in H_2_O once weekly i.p.; cariporide 3 mg/kg in 50:1 PBS:DMSO once daily i.p.; ESL, 175 or 200 mg/kg in 0.5% methylcellulose suspension once daily p.o.) from day 7 post tumour cell implantation. Block randomisation was performed across cage (docetaxel, total 18 mice across six cages, docetaxel n=11, vehicle, n=7; cariporide, total 10 mice across 3 cages, cariporide n=5, vehicle, n=5; ESL, total 7 mice across 2 cages, ESL n=4, vehicle, n=3). Animals in which palpable tumours failed to develop were excluded. Drug stocks were prepared as follows: docetaxel, ethanol (final dilution 4% v/v); cariporide, DMSO (final dilution 2%); ESL was directly suspended in vehicle. The randomised treatment group allocation and experimental procedures were performed unblinded; however, blinding was used during image analysis and assessment of outcome measures (^23^Na signal, tumour volume, ADC).

### MRI

All experiments were performed on a 7T preclinical MRI system (Bruker Biospec 70/30 USR AVANCE III, Bruker Biospin GmbH, Ettlingen, D) with a 12 channel RT-shim system (B-S30) and preinstalled 660mT/m imaging gradient set (BGA-12s, Bruker). Data were captured using Paravision 6.0.1 software. ^1^H and ^23^Na MR imaging was achieved using either i) a decoupled ^1^H quadrature volume resonator (Bruker, 300 MHz, 1 kW max, outer diameter 114 mm/inner diameter 72 mm) and bespoke, 3 cm ^23^Na surface coil; or ii) a dual-tuned linear ^1^H/^23^Na volume coil (inner diameter 35 mm, outer diameter 112 mm, coil length 50 mm, RAPID Biomedical GmbH, Rimpar). All live animal experiments were performed with an NaCl phantom (50 mM) to enable normalisation of signal between animals.

Imaging was performed at 2, 3 and 4 weeks (MDA-MB-231 xenografts) or at 7 days (4T1 and EMT6 xenografts) post implant. Mice were anaesthetised (2% isoflurane in 2 l/min O2) and breathing monitored using a pressure sensitive pad (MR-Compatible Model 1025 Monitoring and Gating System, SAII, NY, USA) throughout. Body temperature was maintained at 37°C using a water heated bed. Following ^1^H localiser scans (IntraGate based), all ^23^Na imaging to assess total tissue Na^+^ utilised either a 2D gradient echo Cartesian acquisition sequence (2D Cartesian, ^23^Na surface coil and ^1^H/^23^Na volume coil) or a 3D gradient echo spiral out sequence (3D spiral, ^1^H/^23^Na volume coil only) for readout.

2D Cartesian: TE, 1.512 ms; TR, 50 ms; □, 90°; nominal resolution, 32×32 voxels; FOV, 40×40 mm; 400 averages; 10 minutes 40 seconds acquisition duration.

3D spiral: TE, 1.533 ms; TR, 10 ms; □, 90°; nominal resolution, 50×50×8 voxels; FOV, 30×30×30 mm; 100 averages; 10 minutes 40 seconds acquisition duration, 3D stack of spirals.

DWI was performed with a readout-segmented echo-planar spin echo imaging sequence and Stejskal-Tanner diffusion gradients applied either i) in three perpendicular directions (MDA-MB-231 xenografts, b values (100, 300 and 700 s/mm^2^) or ii) one direction (EMT6, 4T1 allografts, 100 to 800 s/mm^2^ in increments of 100 s/mm^2^); *b* values were selected to maximise the contrast-to-noise ratio between tumour and non-tumour tissue as previously described^87^.

Tumour and phantom regions of interest (ROI) were determined from the ^1^H structural images. To enable the transfer of ROIs, ^23^Na MRI and ^1^H DWI data were matched in position with their respective ^1^H TurboRARE structural images using a bespoke software package developed in MATLAB and ROI data extracted. For data acquired using the ^23^Na surface coil, images were corrected for B1 field inhomogeneity prior to processing using a linear correction derived from a 1 M NaCl phantom. For comparisons in ^23^Na signal, non-tumour ROIs were taken on the contralateral side to the tumour in solid tissue, avoiding abdominal organs. However, movement artefacts caused blurring between phantom and contralateral tissue during some ^1^H DWI recordings; in these instances an ipsilateral non-tumour ROI was selected and where appropriate the corresponding ^23^Na signal extracted.

### Ex vivo SBFI fluorescence imaging

Tumours were isolated immediately following euthanasia and were sliced in chilled phosphate-buffered saline to a thickness of 200 µm using a 5100MZ vibratome (Campden Instruments Ltd). Tumour slices were loaded with SBFI-AM (10 µM) in HEPES-buffered physiological saline solution (HEPES-PSS: 144 mM NaCl, 5.4 mM KCl, 1 mM MgCl_2_, 1.28 mM CaCl_2_, 5 mM HEPES and 5.6 mM glucose, pH 7.2) with 0.08% Pluronic F-127 for two hours at 37 °C, then rinsed and incubated in fresh HEPES-PSS for 40 minutes to allow uncleaved dye to re-equilibrate. Slices were immobilised within a perfusion chamber (RC-26G, Harvard Bioscience) using a slice hold down (SHD-26GH/10, Harvard Bioscience), and were mounted on an imaging system comprised of a TE200 microscope fitted with a Plan Fluor ELWD 20x/0.45 Ph1 objective (Nikon Corporation, Tokyo, Japan) and a Rolera XR 12 Bit Fast 1394 CCD camera (QImaging, Surrey, British Columbia) controlled by SimplePCI software (Hamamatsu). Excitation light (340 and 380 nm, 50 ms exposure) was separated from emitted light using a 400 DCLP dichroic with a D510/80m filter. Na^+^□free PSS was prepared by replacing NaCl with equimolar *N*□methyl□D□glucamine (NMDG, 144 mM). PSS for calibrating intracellular [Na^+^] was prepared containing 10, 20 and 50 mM NaCl, with additional KCl added in place of omitted NaCl: 149.4 mM NaCl + KCl, 1 mM MgCl_2_, 1.28 mM CaCl_2_, 5 mM HEPES, 5.6 mM glucose, adjusted to pH 7.2 with KOH. Following baseline fluorescence recording, cells were perfused with 10 mM [Na^+^]_e_; the Na^+^ ionophores (gramicidin, 3 µM; monensin, 10 µM) were applied to equilibrate [Na^+^]_i_ and [Na^+^]_e_ and a Na^+^/K^+^ ATPase inhibitor (ouabain, 1 mM) was applied to inhibit Na^+^ efflux. [Na^+^]_e_ was sequentially changed to 20 mM and 50 mM (Figure 3Cii)^40^. In each experimental repeat, [Na^+^]_i_ was calibrated using the final three acquisition frames for each perfusion phase. [Na^+^]_i_ was calibrated *in situ* to control for differences between cells^88^.

### Ion-selective microelectrodes

To produce Na^+^ ion-selective microelectrodes (ISMEs), non-filamented borosilicate micropipettes were pulled to a resistance of 5-10 MΩ (P-97 micropipette puller, Sutter Instruments, Novato, CA, USA) and coated with a hydrophobic layer of N,N-dimethyltrimethylsilylamine (Sigma Aldrich, UK). Micropipettes were back-filled with HEPES-PSS (2.5 mM CaCl_2_, otherwise as described above) and front-filled by suction with oil containing Na^+^ ionophores (sodium ionophore II – cocktail A Selectophore™, Sigma Aldrich, UK). An Axon 900B amplifier (Molecular Devices, Sunnyvale, CA) with a 200 GΩ impedance HS-2 headstage was used to measure voltage. Signals were low-pass filtered at 10 Hz and digitised by an NPI LHBF-48X amplifier/filter system (npi electronic GmbH, Bauhofring, Germany). Electrodes were calibrated with HEPES-PSS of variable [Na^+^] (48, 96, 144 and 192 mM, no replacement ion). Isolated tumours were sliced (500 µm) using a vibratome as described above and then maintained at 30 °C at an interface between HEPES-PSS perfusion and humidified air. Recordings were taken from the top surface of tumour slices and from different locations within each slice. ISMEs were calibrated with known [Na^+^] (48, 96, 144 and 192 mM) prior to and after recording. The junction potential offset between the bath and calibration tube was subtracted from the recorded voltage. All recordings were made within 90 minutes of euthanasia.

### Patch-clamp electrophysiology

Plasma membrane Na^+^ currents in cells within acutely isolated tumour slices (200 µm, sliced with a vibratome as described above) were recorded using the patch clamp method in the whole cell configuration, as previously described^37,89^. Borosilicate glass patch pipettes were pulled (P-97 pipette puller, Sutter Instruments) and fire polished to achieve a resistance of 3–5 MΩ when filled with intracellular recording solution (containing 5 mM NaCl, 145 mM CsCl, 2 mM MgCl_2_, 1 mM CaCl_2_, 10 mM HEPES and 11 mM EGTA, (adjusted to pH 7.4 with CsOH) and bathed in HEPES-PSS (2.5 mM CaCl_2_, otherwise as described above). Recordings were performed using a Multiclamp 700B amplifier (Molecular Devices) at room temperature, compensating for series resistance by ≤40%. A Digidata interface (Molecular Devices) was used to digitize currents, which were low pass filtered at 10 kHz and sampled at 50 kHz. All recordings were analysed using pCLAMP 10.7 software (Molecular Devices).

Two voltage clamp protocols were used:

1. To determine the voltage-dependence of activation, cells were held at -120 mV for 250 ms and then depolarised to between -85 mV and +30 mV for 50 ms (5 mV steps).
2. To determine the voltage-dependence of steady-state inactivation, cells were held at -120 mV for 250 ms, and then sequentially held at a prepulse potential between -120 mV and +30 mV for 250 ms (5 mV steps) followed by a test pulse to -10 mV for 50 ms.

### Inductively-coupled plasma mass spectrometry (ICP-MS)

Total tissue [Na^+^] was determined by inductively coupled plasma mass spectrometry (ICP-MS, 7700x, Agilent). Before digestion, tumour and healthy mammary gland samples were freeze dried overnight (PowerDry LL1500) and both wet and dry weights recorded. The dried sample was digested with nitric acid and hydrogen peroxide (4:1, trace metal grade) within a PTFE digestion vessel using an ETHOS UP microwave digestion system (Milstone, Bergamo, Italy) and the digestion method “SK-CL-002-Animal Tissue”. Digestate was diluted to volume using purified H_2_O. Trace metal contamination from glassware was eliminated using a traceClean acid-steam cleaning system (Milestone). Na^+^ content was quantified using certified reference standards (multi-element environmental calibration standard, Agilent, 5183-4688) and a 10 ppb internal standard solution prepared from a certified reference solution (Agilent, 5188-6526). To determine [Na^+^], the weight (g) of Na^+^ per kg of fresh weight tumour was calculated from the parts per thousand Na^+^ concentration of the dry sample multiplied by the dry sample weight to wet sample weight ratio. This value was then divided by the MW of Na^+^ to give [Na^+^] in mol/kg (reported as mmol/kg).

### LC-MS analysis of plasma and tumour [licarbazepine]

Tissue samples from xenograft tumours were snap frozen and homogenised in chilled 0.1 M PB (pH 5, 1:4 w/v). Lysates were centrifuged (4800 rpm, 4□C, 15 min) and protein was precipitated from the supernatant by mixing lysate with acetonitrile (1:20) containing internal standard (10,11-dihydrocarbamazepine, 10.5 ng/ml, Sigma Aldrich, UK), vortexing, and incubating at -20□C for 30 min. Precipitated protein was removed (centrifuged at 10000 rcf for 7 min at 4□C) and the supernatant analysed by LC-MS (injection volume: 2 µL). Using a 6-level calibration curve of licarbazepine (3.7–900 ng/ml, Tocris Bioscience), normalised data was quantified using a quadratic equation fit (R^2^ = 1.0). Limit of detection and limit of quantification were 0.64 and 1.91 ng/mL, respectively. The LC-MS system consisted of an Acquity I-Class LC (Waters, Elstree, UK) and a Waters BEH C18 column (50 x 2.1 mm, 1.7 µm) including a filter frit, maintained at 40□C. Sample temperature was 10□C). Mobile phase consisted of A) 10% (v/v) acetonitrile with 0.1% acetic acid, and B) 90% acetonitrile with 0.1% acetic acid. LC-MS gradient was 0% B 0 min, 40% B 3 min; 100% B 3.1 -5.5 min, 0% B 5.6–7 min. The LC was connected to a Synapt G2-Si (Waters, Elstree, UK) operating in positive electrospray ionisation sensitivity mode (capillary voltage, 0.5 kV; sampling cone voltage, 40 V; source and desolvation temperatures, 150□ and 500□C; cone gas, 10 L/h; desolvation gas, 1000 L/h; nebuliser, 6.5 bar). Precursor/fragment pairs in the multiple reaction monitoring (MRM) scans were 237.12/194.10 for LIC, 239.13/194.10 for DHC. Data were analysed using Skyline 20.2.0^90^. Tumour and plasma [licarbazepine], were reported as ng licarbazepine per mg tumour (ng/mg) and µM, respectively.

### Immunohistochemistry

Isolated tumour samples were cryoprotected in sequential overnight incubations in 4% paraformaldehyde, 10% sucrose then 30% sucrose before embedding in frozen optimal cutting temperature embedding medium (VWR International). Samples were sectioned (12 µm thickness) onto SuperFrost Plus™ Adhesion slides (Thermo Scientific, Waltham, MA) using a LEICA CM1950 cryostat. Immunohistochemistry was performed using a rabbit anti-Ki67 primary antibody (1:500; Abcam) with an Alexa-568-conjugated goat anti-rabbit secondary antibody (1:500; Invitrogen). Sections were mounted in Prolong Gold with DAPI (Invitrogen) and scanned using a Zeiss AxioScan.Z1 slide scanner (20x magnification).

### Data analysis

All MRI image processing was performed using MATLAB R2019b (MathWorks, Natick, MA). For ^23^Na MRI, mean and maximum signal data were normalised to that of the phantom (50 mM NaCl) so that they represented a Na^+^ concentration measurement. Normal distribution of data was confirmed using a Shapiro-Wilk test. Statistical comparisons performed at each independent timepoint using an unpaired, two way Student’s t-test. Correlations were assessed using a Pearson’s r test. Comparisons between tumour models (Figure 6) were performed using a one-way ANOVA with post-hoc Tukey test for multiple comparisons.

DWI data were analysed using RStudio version 1.2.5033 (RStudio Inc., Boston, MA). ADC was calculated as ADC = −ln(S/S_0_)/b, where b is the b value in seconds/mm^2^ and S_0_ is the signal intensity with no diffusion gradients. Principal component analysis was performed using the prcomp() function and the “factoextra” package (fviz_pca_biplot(), fviz_pca_ind()).

Linear discriminant analysis (LDA) with leave-one-out cross validation (LOOCV) were performed using the “MASS” package (lda()). Due to the limited size of the training data sets in Figure 2, LOOCV was used as an initial estimate of model classification accuracy, while an independent test set (control group from Figure 4) was used to compare the prediction accuracy of models trained on maximum ^23^Na signal vs ADC. Receiver operating characteristic curve was determined using the “ROCR” package (prediction(), performance()).

To calibrate [Na^+^]_e_, ISME recordings from standards were fitted to a non-linear regression (Padé (1,1,) approximant), and tumour slice recordings interpolated. Similarly, SBFI fluorescence recordings for each calibration step were fit to nonlinear regression (n=3 experiments, one-phase association) and resting [Na^+^]_i_ (baseline fluorescence) interpolated. Quantification of Ki67-positive nuclei (as a percentage of nuclei count, DAPI) was performed using ZEN 3.2 (Zeiss, Oberkochen, Germany) and ImageJ 1.53c (NIH, public domain) particle count (50 pixel minimum size) with the classic watershed method applied.

The voltage-dependence of activation and steady-state inactivation were determined as follows:

1. Voltage-dependence of activation: G = I/(V_m_ – V_rev_), where G is conductance, I is current, V_m_ is the membrane voltage and V_rev_ is the reversal potential for Na^+^ (+84.64 mV, derived from the Nernst equation). Conductance was normalised G/G_max_.
2. Voltage-dependence of steady-state inactivation: Current was normalised I/I_max_.

Both data sets were fitted to Boltzmann sigmoidal curves. The resulting V_1/2_ + k values (individual cells) were compared between groups using an unpaired, two-way Student’s t-test.

All statistical comparisons were performed using Graphpad Prism 8.3.1 (GraphPad Software, San Diego, CA).

## Acknowledgements

This work was supported by Cancer Research UK (A25922), Breast Cancer Now (2015NovPhD572) and EPSRC Impact Accelerator Award. We thank the Biorenewables Development Centre (Dunnington, York) for their support with the tumour sample ICP-MS analysis, and the University of York Bioscience Technology Facility for their support with tumour and blood plasma LC-MS analysis.

## Author contributions

The project was designed by W.J.B., A.J.K., and A.D.J.; experiments were carried out by A.D.J., T.K.L, S.L., M.C.L., A.J.K.; ^23^Na coil design and setup of imaging protocols were carried out by A.J.K., J.D.K., F.R. and M.A.M.; Data analysis was performed by A.D.J., T.K.L, A.J.K., S.L., L.P., L.W., J.M.O.; the manuscript was prepared by A.D.J., T.K.L., W.J.B., A.J.K.; A.D.J., T.K.L., W.J.B., A.J.K., J.D.K., F.J.G., L.W., M.A.M., and G.B. contributed to the interpretation of the data and critical revision of the paper for important intellectual content.

