## Supplementary Figures for "Sodium accumulation in breast cancer predicts malignancy and treatment response"

[1]Department of Biology, University of York. [2]York Biomedical Research Institute, University of York. [3]Department of Radiology & NIHR Cambridge Biomedical Research Centre, University of Cambridge. [4]Department of Mathematics, University of York. [5]Department of Chemistry, University of York. [6]Bioscience Technology Facility, Department of Biology, University of York. [7]Department of Physics, University of York. [8]Mohn Medical Imaging and Visualization Centre, Haukeland University Hospital Bergen.

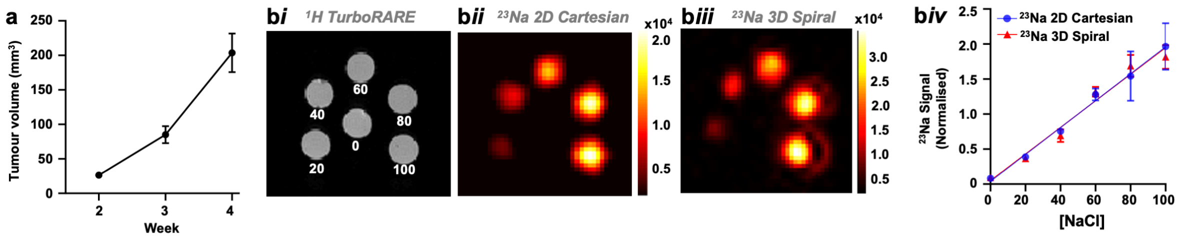


**Supplementary Figure 1**: (***a***) Growth rate of orthotopic MDA-MB-231 tumour xenografts; cells seeded at 5 x $10^5$ cells per mouse, 50:50 PBS:Matrigel (n=6, volume measured by multislice (1 mm) $^{1}$H TurboRARE imaging at Weeks 2, 3 and 4 post-implant). (***bi***) Axial slice of six 4 mm diameter phantoms containing known NaCl concentrations (1% agarose, [NaCl] in mM annotated) and corresponding representative $^{23}$Na images using $^{23}$Na FLASH (**b*ii***) and $^{23}$Na spiral (**b*iii***) acquisition. (**b*iv***) $^{23}$Na signal (normalised to that of 50 mM NaCl) was linear with [NaCl] for both the $^{23}$Na FLASH (blue) and $^{23}$Na spiral (red) acquisition ($^{23}$Na FLASH: $R^2$ = 0.7856; 0 mM and, 100 mM, n=3 pixels; 20–80 mM, n=4 pixels. $^{23}$Na spiral: $R^2$ = 0.8565; 0 mM, n=8 pixels; 20–80 mM, n=4 pixels; 100 mM, n=6 pixels). Data represent group mean ± SEM.

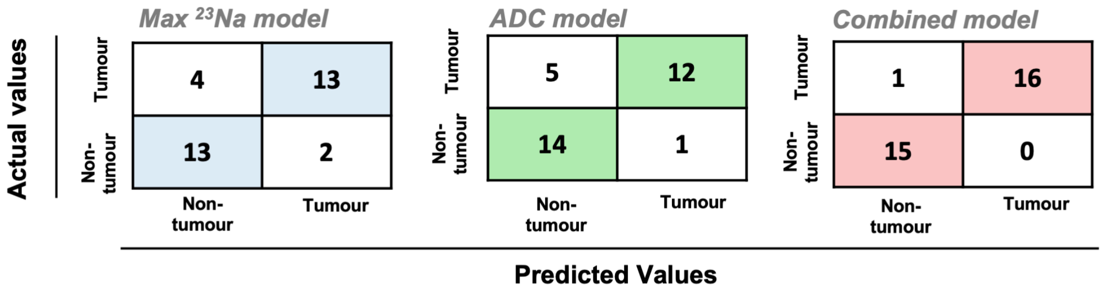


**Supplementary Figure 2**: **Confusion matrices for sensitivity and specificity values presented in Figure 2G.** Linear discriminant analysis models were trained on maximum $^{23}$Na values alone, ADC values alone, or a combination of both parameters, using region of interest (ROI) data from tumour and non-tumour regions (data from Figure Ei, n=30 regions from all timepoints). These models were then used to classify regions from the independent docetaxel cohort vehicle group (data from Figure 5, n=32 regions from all timepoints).

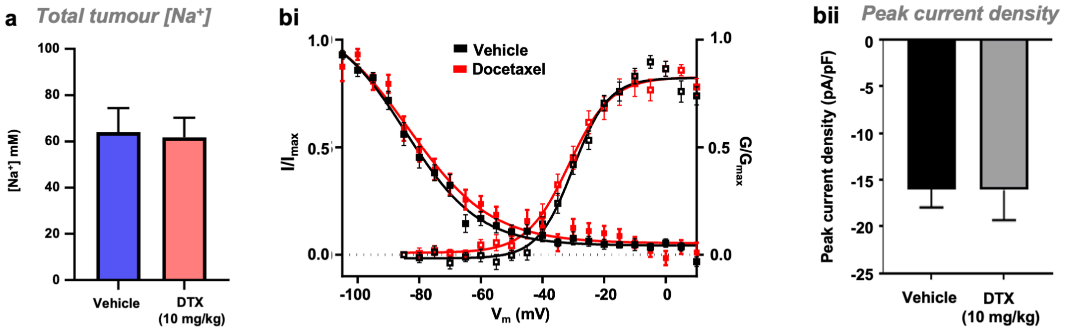


**Supplementary Figure 3: Docetaxel has no effect on voltage gated $Na^+$ channel current in acutely isolated tumour slices** (**a**) Total tumour [$Na^+$] from acutely isolated (Week 4) MDA-MB-231 tumour xenograft samples treated with either docetaxel (10 mg/kg once weekly i.p. From day 7, n=3) or vehicle (n=4), as determined by inductively-coupled plasma mass spectrometry. (**b**) Activation and steady-state inactivation (SSI) were determined for voltage-gated $Na^+$ channels within intact MDA-MB-231 tumour slices from animals treated with docetaxel (red squares, 10 mg/kg i.p. once weekly beginning day 7 post implant) or vehicle (black squares). Tumour slices (200 μm) were isolated following Week 4 MRI. To determine SSI, a holding potential of -120 mV was applied, followed sequentially by 250 ms conditioning voltage prepulses between -105 and +30 mV (5 mV steps) and 50 ms test pulses at -10 mV; the resulting normalized currents ($I/I_{max}$) were plotted as a function of the prepulse voltage (docetaxel, n=14; vehicle, n=23).To determine activation, a holding potential of -120 mV was applied, and currents elicited by 5 mV depolarising steps from -85 to +30 mV for 30 ms; normalized conductance ($G/G_{max}$) was calculated from current and plotted as a function of voltage (docetaxel, n=15; vehicle, n=24). Both activation and SSI curves fitted with Boltzmann functions. (**c**) Quantified peak current density recorded in docetaxel (n=15) and vehicle-treated (n=24) tumour slices. Data represent mean ± SEM.

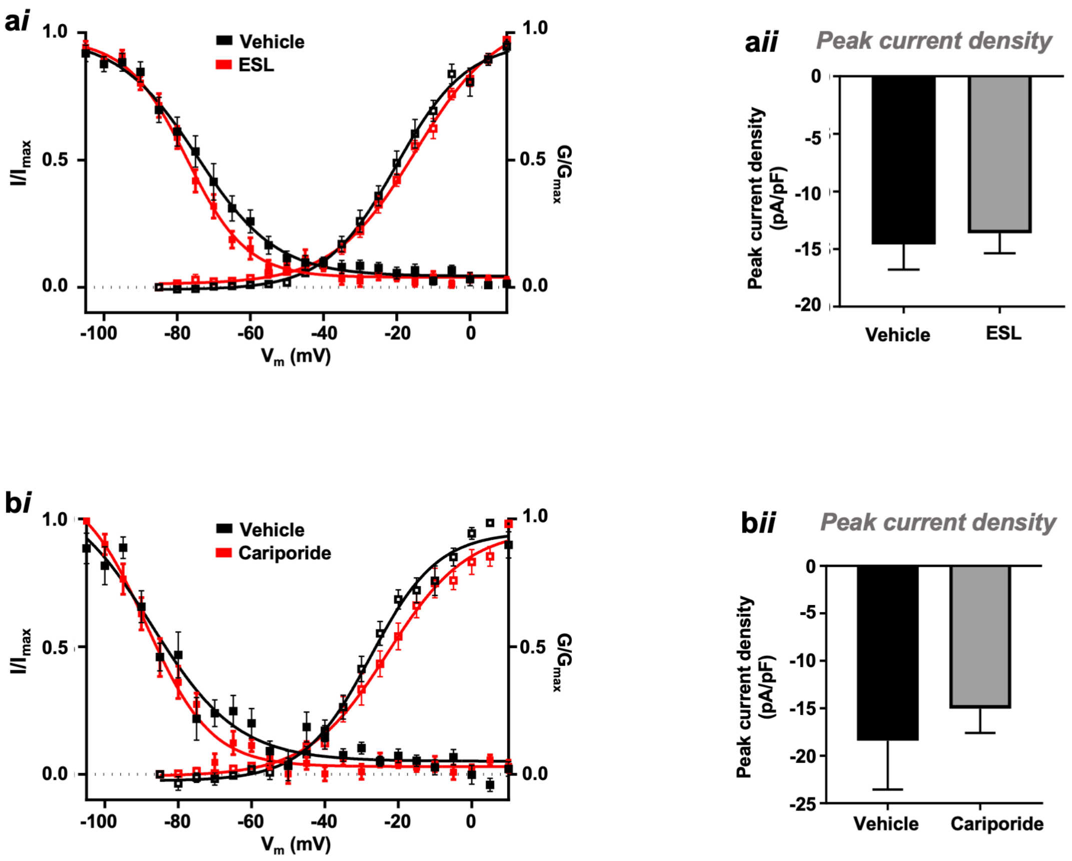


**Supplementary Figure 4. Eslicarbazepine acetate (ESL) and cariporide have no effect on voltage gated $Na^+$ channel current in acutely isolated tumour slices.** Activation and steady-state inactivation (SSI) were determined for voltage-gated $Na^+$ channels within intact MDA-MB-231 tumour slices from animals treated with either eslicarbazepine acetate vs vehicle (***ai***, red squares, 200 mg/kg p.o. once daily (SSI, n=17; activation n=20) beginning day 7 post implant; vehicle (SSI, n=14; activation, n=16) shown in black squares) or cariporide (***bi***, red squares, 3 mg/kg i.p. once daily (SSI and activation, n=11) beginning day 7 post implant; vehicle (SSI and activation, n=9) shown in black squares). Tumour slices (200 μm) were isolated following Week 4 MRI. To determine SSI, a holding potential of -120 mV was applied, followed sequentially by 250 ms conditioning voltage prepulses between -105 and +30 mV (5 mV steps) and 50 ms test pulses at -10 mV; the resulting normalized currents ($I/I_{max}$) were plotted as a function of the prepulse voltage (docetaxel, n=14; vehicle, n=23).To determine activation, a holding potential of -120 mV was applied, and currents elicited by 5 mV depolarising steps from -85 to +30 mV for 30 ms; normalized conductance ($G/G_{max}$) was calculated from current and plotted as a function of voltage (docetaxel, n=15; vehicle, n=24). Both activation and SSI curves fitted with Boltzmann functions. Peak current densities were quantified in slices from tumours treated with either (***aii***) ESL (n=20) or vehicle (n=16) or (***bii***) cariporide (n=11) or vehicle (n=9). Data represent mean ± SEM.
